# Bioactive Cationic Lipidated Oligomers (CLOs) as Antimicrobial Materials: Metabolomic Insights into MRSA Membrane Disruption

**DOI:** 10.64898/2025.12.03.692207

**Authors:** Muhammad Bilal Hassan Mahboob, Jessica R. Tait, Dovile Anderson, Maytham Hussein, John F. Quinn, Tony Velkov, Darren. J. Creek, Michael R. Whittaker, Cornelia B. Landersdorfer

## Abstract

The escalating incidence of antimicrobial resistance in *Staphylococcus aureus*, particularly the methicillin-resistant strain (MRSA), necessitates the development of novel therapeutic strategies. Cationic lipidated oligomers (CLOs) have emerged as promising membrane-active antimicrobial agents; however, their mechanisms of action remain insufficiently understood. In this study, untargeted metabolomics was employed to systematically profile the temporal metabolic perturbations induced by two structurally distinct CLOs, C_12_-o-DMEN-10 and C_12_-o-BEDA-10, in MRSA across four defined time points (0.25, 0.5, 1, and 3 hours). These CLOs previously demonstrated differential antibacterial activity, as evidenced by dose-dependent propidium iodide (PI) uptake and growth inhibition assays. Metabolomic analysis revealed pronounced and sustained disruptions in bacterial membrane lipid metabolism, including significant depletion of phosphatidylglycerols (≥ −2.5 log_2_FC, *p* < 0.05), alongside elevated levels of phosphatidylethanolamines, lysophospholipids, and fatty acid-derived metabolites indicative of membrane destabilization and lipid remodelling. Although both CLOs affected overlapping metabolic pathways, they differed in the extent and temporal dynamics of their effects. These findings provide mechanistic insights into CLO-mediated antibacterial activity and highlight the value of metabolomics in elucidating both direct and downstream cellular responses, which extend beyond the scope of conventional membrane integrity assays, such as PI fluorescence.

## Introduction

Antimicrobial resistance (AMR) represents one of the most urgent global health threats of the 21^st^ century [1, 2]. The World Health Organization (WHO) has identified AMR as a critical disruptor to the efficacy of modern antimicrobial medicines [3, 4]. In 2019 alone, bacterial AMR was directly responsible for approximately 1.27 million deaths and contributed to nearly 4.95 million deaths globally [5]. Projections estimate that without immediate and coordinated action, AMR could cause up to 10 million deaths annually by 2050, accompanied by substantial economic losses, including an estimated additional US$1 trillion in healthcare costs and US$1– 3.4 trillion in annual global GDP losses by 2030 [6-9]. The primary drivers of AMR include the misuse and overuse of antimicrobial agents across human, animal, and agricultural sectors, compounded by socioeconomic inequalities, especially in low- and middle-income countries [10, 11]. The delay in the antibiotic development pipeline, failure to identify antibiotics with new mechanisms of action, and resistance against existing antimicrobials further intensify the crisis [12, 13]. In the absence of effective antibiotics, routine medical procedures such as surgeries, organ transplants, and cancer chemotherapies become significantly riskier, underlining the urgent need for novel therapeutic strategies [14-16].

Among the resistant pathogens contributing to this global burden, *Staphylococcus aureus* (*S. aureus*), particularly its methicillin-resistant form i.e., methicillin-resistant *S. aureus* (MRSA), has been declared as a high-priority threat by WHO [17-19]. MRSA is one of the leading causes of healthcare-associated infections worldwide and has been categorized as a “serious threat” by the U.S. Centers for Disease Control and Prevention (CDC) [18, 20, 21]. In 2019, MRSA was the deadliest drug-resistant pathogen globally, responsible for an estimated 121,000 deaths attributable to AMR, according to the Institute for Health Metrics and Evaluation (IHME) [22]. The alarming increase in resistance among *S. aureus* strains highlights the urgent necessity for developing alternative antimicrobial agents with novel modes of action [23-27].

In recent years, antimicrobial polymers have gained attention as promising candidates to counter multidrug-resistant (MDR) pathogens [28-30]. These materials have co-opted the structural elements in natural antimicrobial peptides: cationic membrane binding and lipid membrane-disrupting elements [31]. Among these, cationic lipidated oligomers (CLOs) are synthetic amphiphilic macromolecules designed to mimic the structural and functional properties of antimicrobial peptides (AMPs) more closely [32, 33]. CLOs consist of cationic groups distributed along short polymeric chains, which facilitate electrostatic interactions with negatively charged bacterial membranes, and terminal hydrophobic lipid tails, which insert into and disrupt the membrane, leading to cell lysis [34, 35]. Their tuneable amphiphilic nature enables them to exert potent antimicrobial effects across a range of priority bacterial and fungal pathogens [36, 37]. Moreover, CLOs offer significant advantages over natural AMPs, including enhanced stability, facile synthesis, access to a wider range of cationic and lipid functionalities, and therefore a greater structure-activity relationship space [37]. However, despite their promising antibacterial activities, the detailed mechanisms by which CLOs exert microbial killing remain poorly understood [38].

To address this knowledge gap we have employed untargeted metabolomics, which is a powerful tool for elucidating phenotypic and biochemical changes in microbial cells upon treatment with antimicrobial agents [39, 40]. Metabolomics, particularly untargeted approaches utilizing liquid chromatography mass spectrometry (LC-MS), has emerged as a robust analytical strategy for detecting nuanced metabolic alterations across biological systems [41, 42]. These techniques provide mechanistic insights that are instrumental in guiding the structural refinement of lead compounds, informing rational design of combination therapies, and facilitating the monitoring of antimicrobial resistance in clinical settings [43, 44]. Our previously published work, Hussein *et al*. (2024), remains the only study to date employing metabolomics to elucidate the antimicrobial mechanism of antimicrobial polymers, including CLOs [45].

In the present study, we conducted a comprehensive untargeted metabolomics investigation to elucidate the antimicrobial mechanism of action(s) of two structurally distinct CLOs, each comprising a dodecyl (C_12_) lipid tail, but with differing cationic functionality (tertiary or primary amine) against MRSA ATCC 43300. We delineated the principal metabolic pathways disrupted by CLO exposure, thereby advancing a mechanistic understanding of the cellular biochemical perturbations that underpin the structure-property relationships governing CLO-mediated bactericidal activity. These insights provide a critical framework for the rational design and development of next-generation antimicrobial agents.

## Results and discussions

The synthesis, characterisation and toxicity profile of the CLOs [C_12_-o-DMEN-10 (tertiary amine cationicity) and C_12_-o-BEDA-10 (primary amine cationicity)] have been previously published [46]. In this aforementioned published work, the antibacterial efficacy of the CLOs was evaluated against MRSA ATCC 43300, revealing minimum inhibitory concentrations (MICs) of 128 µg/mL and 64 µg/mL, respectively [46]. Furthermore, these CLOs demonstrated distinct mechanistic profiles as evidenced by a comparative assessment of membrane disruption using a dose-dependent propidium iodide (PI) fluorescence assay and bacterial growth inhibition **(Fig. S1)**. As illustrated in **Fig. S1(a)**, CLO [C_12_-o-DMEN-10, bearing tertiary amine functionality] induced over 50% propidium iodide (PI) fluorescence at its MIC, i.e., 128 µg/mL. This substantial membrane permeability suggested that C_12_-o-DMEN-10 exerts its antibacterial effect primarily through membrane disruption. Conversely, the CLO [C_12_-o-BEDA-10, featuring primary amine cationicity] demonstrated negligible PI fluorescence across the tested concentration range as depicted in **Fig. S1(b)**, indicating minimal membrane perturbation and suggesting an alternative, non-membranolytic mechanism of bacterial killing [46]. Collectively, these findings highlight that the divergent antibacterial activities of the CLOs are likely driven by differences in their structural features, specifically cationic functionalities (tertiary amine vs. primary amine). Although tertiary versus primary amines were intentionally used as the designed variable, this modification also affects other physicochemical properties such as hydrophobic balance, polarity, protonation behaviour, and overall amphiphilic geometry [47, 48]. These parameters can contribute collectively to the different magnitudes and temporal profiles of metabolic disruption observed between these two CLOs. To further elucidate these mechanistic distinctions, an untargeted metabolomics investigation was performed, using an initial bacterial inoculum of approximately ∼10^8^ colony-forming units (CFU)/mL, with samples collected at 0.25 h, 0.5 h, 1 h, and 3 h post-treatment. A static concentration time-kill assay was initially performed to guide the selection of both appropriate time points and CLO concentrations for the metabolomics experiment. Based on the observed bacterial killing kinetics, four time points, 0.25, 0.5, 1, and 3 h post-treatment, were selected to capture the dynamic progression of metabolic perturbations. CLOs were administered at concentrations that elicited bactericidal activity, specifically at 2x MIC for C_12_-o-DMEN-10 and 4x MIC for C_12_-o-BEDA-10, to ensure the induction of robust intracellular responses i.e., approximately ∼1.5 log_10_ CFU/mL relative to untreated controls **(Fig. 1)**. These parameters were chosen to enable high-resolution temporal profiling of CLO-induced biochemical alterations during bacterial killing. The metabolomic alterations and disrupted metabolic pathways in MRSA following CLO exposure are discussed in detail below.

**Figure 1.**
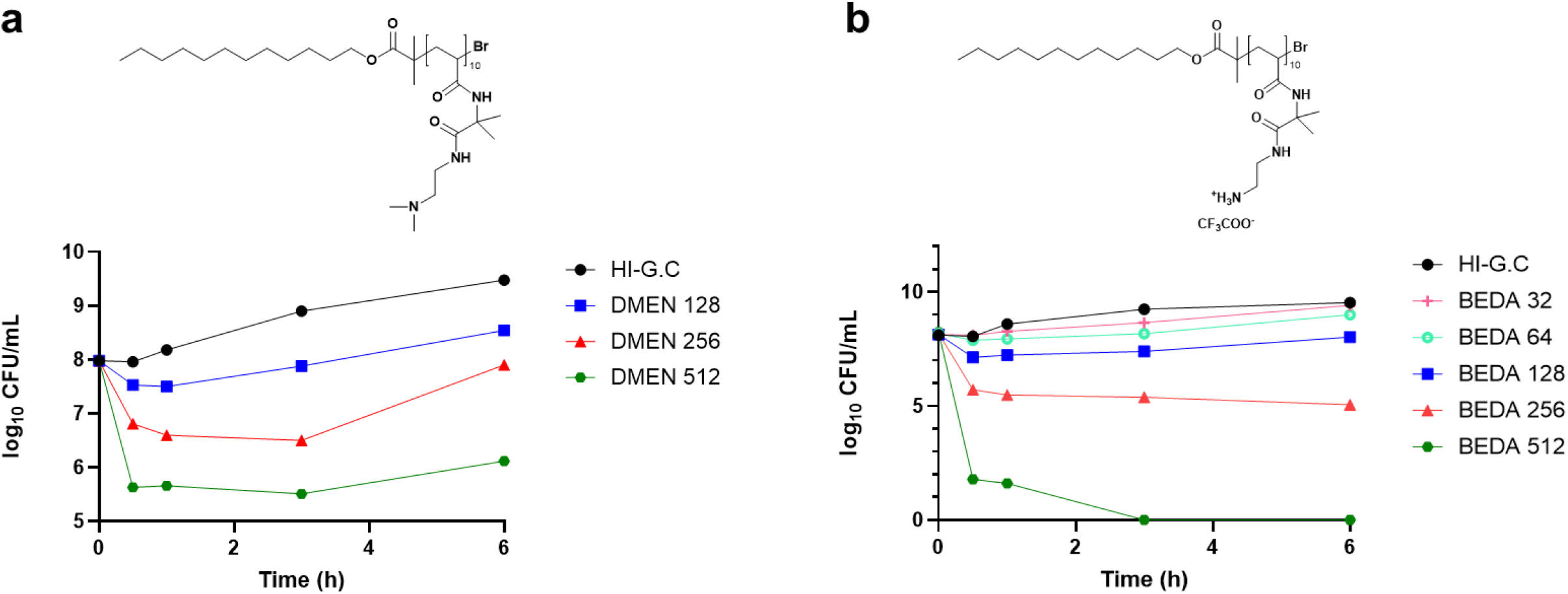
Killing kinetics of MRSA ATCC 43300 (initial inoculum∼10^8^ log_10_ CFU/mL), after treatment with different concentrations of CLOs [μg/mL] **(a)** C_12_-o-DMEN-10 (128, 256, 512 μg/mL) and **(b)** C_12_-o-BEDA-10 (32, 64,128, 256, 512 μg/mL)] (n=1).

### Global metabolic disruptions in MRSA ATCC 43300 in response to CLOs

A total of 1,018 putative metabolites were identified across both treatments, i.e., C_12_-o-DMEN-10 and C_12_-o-BEDA-10. Metabolite annotation was performed using established databases, including BioCyc, KEGG, HMDB and LipidMaps, which also facilitated the mapping of putative metabolites to pathways. Principal component analysis (PCA) demonstrated clear separation between treated and control groups at all time points, indicating substantial CLO-induced metabolic perturbation in MRSA **(Fig. S2)**. In line with PCA, heatmaps further revealed dynamic fluctuations in metabolite intensities following treatment with CLOs **(Fig. 2)**. Univariate statistical analysis was conducted [using two-sample t-tests with a log_2_fold-change (FC) threshold ≥ 2 (corresponding to a ∼4-fold change in abundance) and a false discovery rate (FDR)-adjusted *p*-value < 0.05] to determine significantly perturbed metabolites at 0.25 h, 0.5 h, 1 h, and 3 h post-treatment **(Table S1)**. This analysis revealed approximately 203 and 198 significantly perturbed metabolites following challenge with C_12_-o-DMEN-10 and C_12_-o-BEDA-10, respectively. Specifically, C_12_-o-DMEN-10 perturbed 177, 185, 183, and 177 metabolites at 0.25 h, 0.5 h, 1 h, and 3 h, while for C_12_-o-BEDA-10, numbers were 155, 178, 167, and 166, respectively **(Fig. S3)**. Only a few metabolites were uniquely altered at individual time points, suggesting that both CLOs triggered a sustained and consistent metabolic response **(Fig. S3)**. The classification of significantly perturbed metabolites demonstrated that a wide range of pathways representing carbohydrates, amino acids, nucleotides, and peptides were affected, with most metabolite levels being depleted after treatment. As these pathways are predominantly comprised of polar metabolites, it is assumed that the drastic depletion of most metabolites observed at all time-points (shown in **Fig. 2**) is due to membrane disruption and leakage of intracellular metabolites. Interestingly, lipids and fatty acids exhibited the substantial and consistently sustained alterations that included some lipids decreasing, while others increased in abundance, indicating that cell membranes were retained, and that these CLOs induced specific changes in lipid metabolism and membrane composition **(Table S1)**.

**Figure 2.**
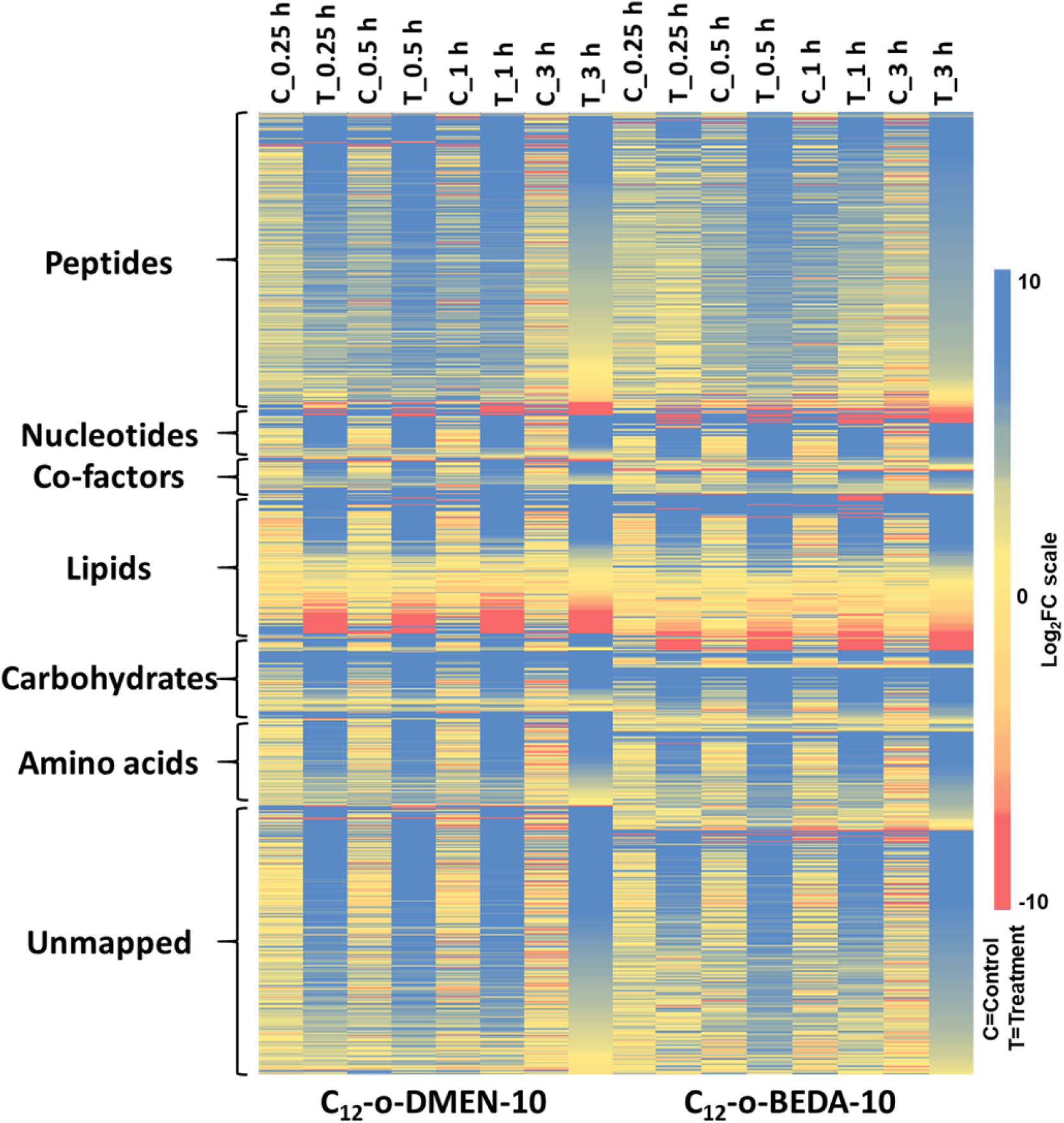
Heatmaps by treatment with 256 µg/mL of CLOs [C_12_-o-DMEN-10 and C_12_-o-BEDA-10] at 0.25 h, 0.5 h, 1 h and 3 h (T_0.25 h, T_0.5 h, T_1 h and T_3 h, respectively) and control (C_ 0.25 h, C_ 0.5 h, C_1 h and C_3 h) in strain MRSA ATCC 43300 using untargeted metabolomics.

### Perturbations in the lipids (glycerophospholipids and lysophospholipids) metabolism

Lipids, including glycerophospholipids, are critical components of the cytoplasmic membrane in Gram-positive bacteria such as *S. aureus*, contributing to membrane integrity, fluidity, and adaptive responses to environmental and antimicrobial stress [49-52]. In *S. aureus*, the major glycerophospholipids include phosphatidylglycerol (PG, ∼50–60%), phosphatidylethanolamine (PE, ∼20–30%), and smaller amounts of phosphatidic acid (PA, ∼1– 5%), phosphatidylserine (PS), and phosphatidylinositol (PI), each comprising less than 5% [53, 54]. A major resistance mechanism in MRSA involves the conversion of PG to lysyl-phosphatidylglycerol (L-PG, ∼10–15%), which increases the membrane’s positive charge, thereby repelling cationic antimicrobial peptides and antibiotics from bacterial action [55-59].

Additionally, lysophospholipids, including Lyso-PG, generated through membrane remodeling, contribute to stress adaptation [60]. Given their central roles in membrane function and resistance, targeting these lipid biosynthetic and modification pathways offers a promising strategy for the development of next-generation antimicrobials against MRSA [61, 62]. Both CLOs caused significant perturbations in the lipid metabolism of MRSA across all timepoints.

At 0.25 h post-treatment, both C_12_-o-DMEN-10 and C_12_-o-BEDA-10 induced rapid and significant alterations in the MRSA lipid profile, particularly within the glycerophospholipids. C_12_-o-DMEN-10 elicited a more pronounced response, with marked depletion in PGs including PG(34:0), PG(35:0), and PG(33:0) (≥−6.5-log_2_FC, *p* < 0.05). In comparison, C_12_-o-BEDA-10 produced milder, yet notable, reductions in the same lipids (≥−5.0-log_2_FC, *p* < 0.05). Concurrently, an accumulation of lysophospholipids, especially LysoPE(18:1) and LysoPE(19:1), was observed under both treatments, with greater elevations under C_12_-o-DMEN-10 treatment [C_12_-o-DMEN-10 (≥−1.5-log_2_FC, *p* < 0.05) than C_12_-o-BEDA-10 (≥−1.0-log_2_FC, *p* < 0.05)]. These early changes suggest rapid membrane perturbation and enhanced phospholipid turnover, reflective of immediate stress responses involving membrane degradation and remodeling [63, 64] **(Fig. 3 a,b)**.

**Figure 3.**
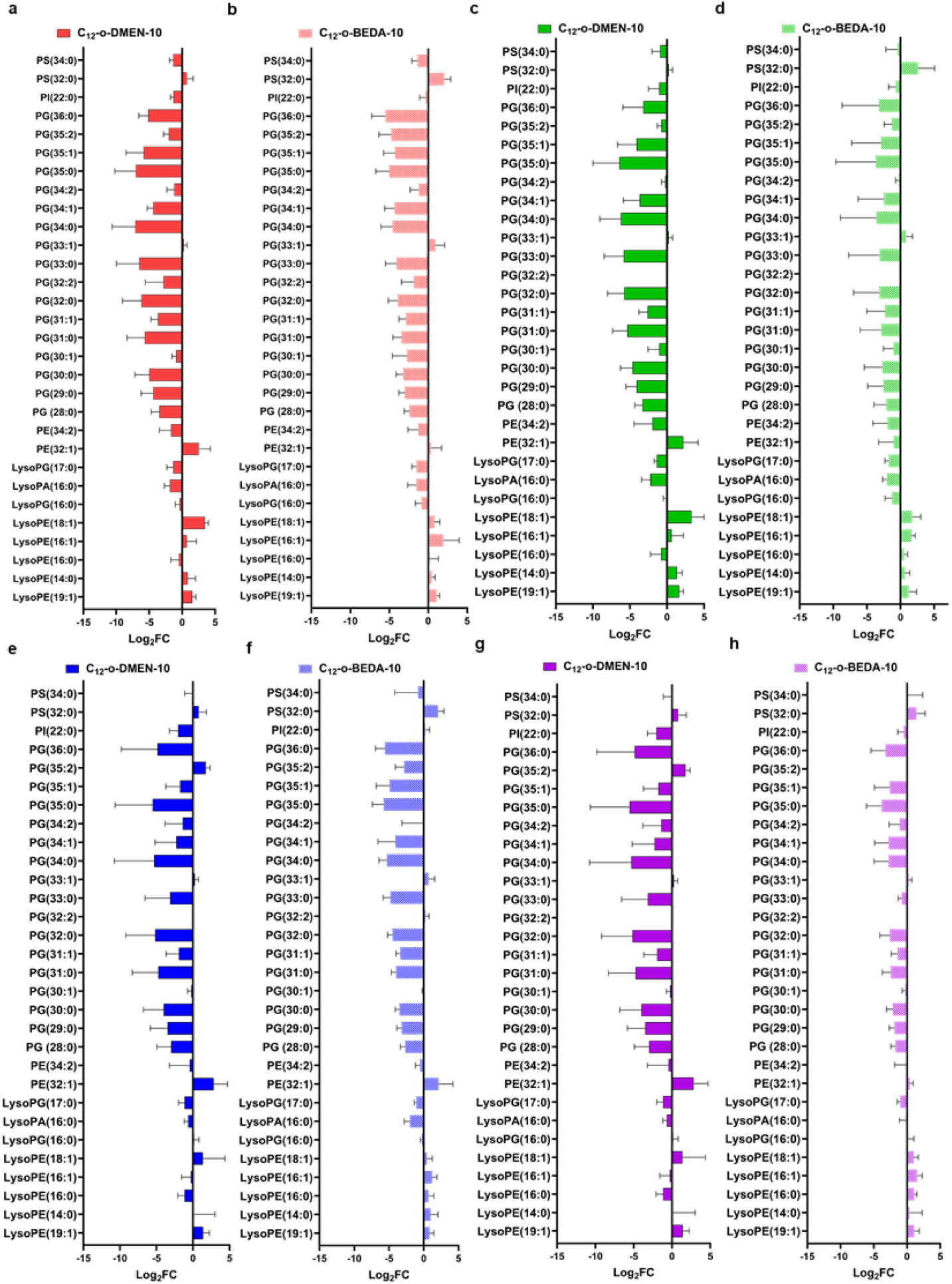
Significantly perturbed lipids in MRSA ATCC 43300 following treatments with CLOs [C_12_-o-DMEN-10 (solid colours) and C_12_-o-BEDA-10 (dotted colour)] at 0.25 h (**a**,**b**; red), 0.5 h (**c**,**d**; green), 1 h (**e**,**f**; blue), 3 h (**g**,**h**; purple). Lipid names are putatively assigned based on accurate mass (≥1.0-log_2_-FC; *p* < 0.05). PE, phosphoethanolamines; PG, glycerophosphoglycerols; PS, glycerophosphoserines; PA, glycerophosphates; LysoPE, lysophosphatidylethanolamines; LysoPG, lysophosphatidylglycerols.

At 0.5 h, the lipid alterations intensified under both treatments, with continued depletion of key PG lipids. C_12_-o-DMEN-10 maintained a greater impact, with PG(35:0), PG(34:0), and PG(33:0) showing sustained depletion (≥−5.5-log_2_FC, *p* < 0.05). C_12_-o-BEDA-10 also affected PGs, however, the effect was generally less pronounced (≥−3.5-log_2_FC, *p* < 0.05). An increase in PE(32:1) was particularly evident with C_12_-o-DMEN-10, indicating a potential shift in lipid composition to maintain membrane integrity [65]. Lysophospholipid levels, including LysoPE(18:1) and LysoPE(19:1), remained elevated under both CLOs treatment, further supporting ongoing membrane lipid remodelling processes in response to CLO exposure [66] **(Fig. 3 c,d)**.

At the 1h time point, a broad range of PGs continued to show significant depletion under both treatments. For C_12_-o-DMEN-10, PG(34:0) and PG(35:0) remained highly suppressed (≥−5.53 log_2_FC, and ≥−5.45-log_2_FC, *p* < 0.05 respectively), while other PGs including PG(32:0), PG(33:0), and PG(30:0) also showed substantial reductions (≥−3.0-log_2_FC, *p* < 0.05). C_12_-o-BEDA-10 induced comparable depletions, where the level of PG(34:0), PG(35:0), PG(32:0), and PG(30:0) was further declined (≥−3.0-log_2_FC, *p* < 0.05). Elevation in PE(32:1) was sustained in both treatments (1.97-log_2_FC for C_12_-o-DMEN-10 and 2.15-log_2_FC for C_12_-o-BEDA-10), indicating a persistent shift in membrane lipid composition [67]. Notably, lysophospholipid levels such as LysoPE(18:1) and LysoPE(19:1) declined slightly compared to earlier time points, [LysoPE(18:1) declined from 3.32 to 0.69-log_2_FC under C_12_-o-DMEN-10 and from 1.73 to 0.45-log_2_FC under C_12_-o-BEDA-10, *p* < 0.05], potentially reflecting reintegration into biosynthetic pathways or compensatory membrane repair mechanisms [67, 68] **(Fig. 3 e,f)**.

At 3 h, both CLOs continued to significantly perturb phospholipid metabolism, with C_12_-o-DMEN-10 exerting a more pronounced effect. PGs species remained substantially depleted, particularly under C_12_-o-DMEN-10, where PG(34:0), PG(35:0), and PG(32:0) showed extensive decline (≥−5.0-log_2_FC, *p* < 0.05) compared to milder reductions with C_12_-o-BEDA-10 (≥−2.5-log_2_FC, *p* < 0.05). Lysophospholipids, including LysoPE(18:1) and LysoPE(19:1), remained elevated (≥1.0-log_2_FC, *p* < 0.05), though less pronounced than at earlier time points, indicating ongoing but possibly stabilizing membrane remodelling **(Fig. 3 g,h)**.

Collectively, these findings indicate sustained disruption of lipid homeostasis following exposure to either CLOs, with C_12_-o-DMEN-10 eliciting a more persistent impact.

### Perturbations in fatty acid (FA) metabolism

Fatty acid (FA) metabolism is essential for the growth, membrane integrity, and stress adaptation of Gram-positive bacteria, including methicillin-resistant *S. aureus* (MRSA) [69, 70]. Fatty acids function as essential precursors for the biosynthesis of membrane phospholipids, wherein their saturation levels, chain lengths, and structural modifications critically modulate biophysical membrane properties, i.e., fluidity, permeability, and mechanical stability [71]. Variations in the degree of unsaturation, acyl chain length, and branching significantly influence membrane dynamics, thereby impacting bacterial adaptability and resistance to antimicrobial agents [72]. MRSA’s ability to exploit both endogenous and host-derived fatty acids enhances its metabolic adaptability under antimicrobial stress, positioning fatty acid metabolism as a promising therapeutic target [73]. The study revealed that both CLOs caused significant perturbations in FA metabolism, albeit with differences in magnitude and persistence across all the time points.

At 0.25 h post-treatment, both C_12_-o-DMEN-10 and C_12_-o-BEDA-10 induced pronounced disruptions in FA metabolism, with notable alterations in hydroxylated and dicarboxylic FA. C_12_-o-DMEN-10 resulted in substantial accumulation of hydroxymyristoylcarnitine and hexacosanedioic acid (≥ 5.0-log_2_FC, *p* < 0.05), whereas C_12_-o-BEDA-10 elicited an even greater elevation in hexacosanedioic acid (≥ 10-log_2_FC, *p* < 0.05), indicating a robust and early response to treatment [74]. FA conjugates such as *N*-heptanoylglycine and stearoylglycine were markedly upregulated following C_12_-o-DMEN-10 treatment (≥ 4.0-log_2_FC, *p* < 0.05), but concurrently suppressed under C_12_-o-BEDA-10, suggesting differential regulation of FA metabolism. However, hydroxylated intermediates, including hydroxydocosanoic acid and hydroxy-octadecenoylcarnitine, were consistently elevated (≥ 1.0-log_2_FC, *p* < 0.05) by both CLOs treatments **(Fig. 4)**.

**Figure 4.**
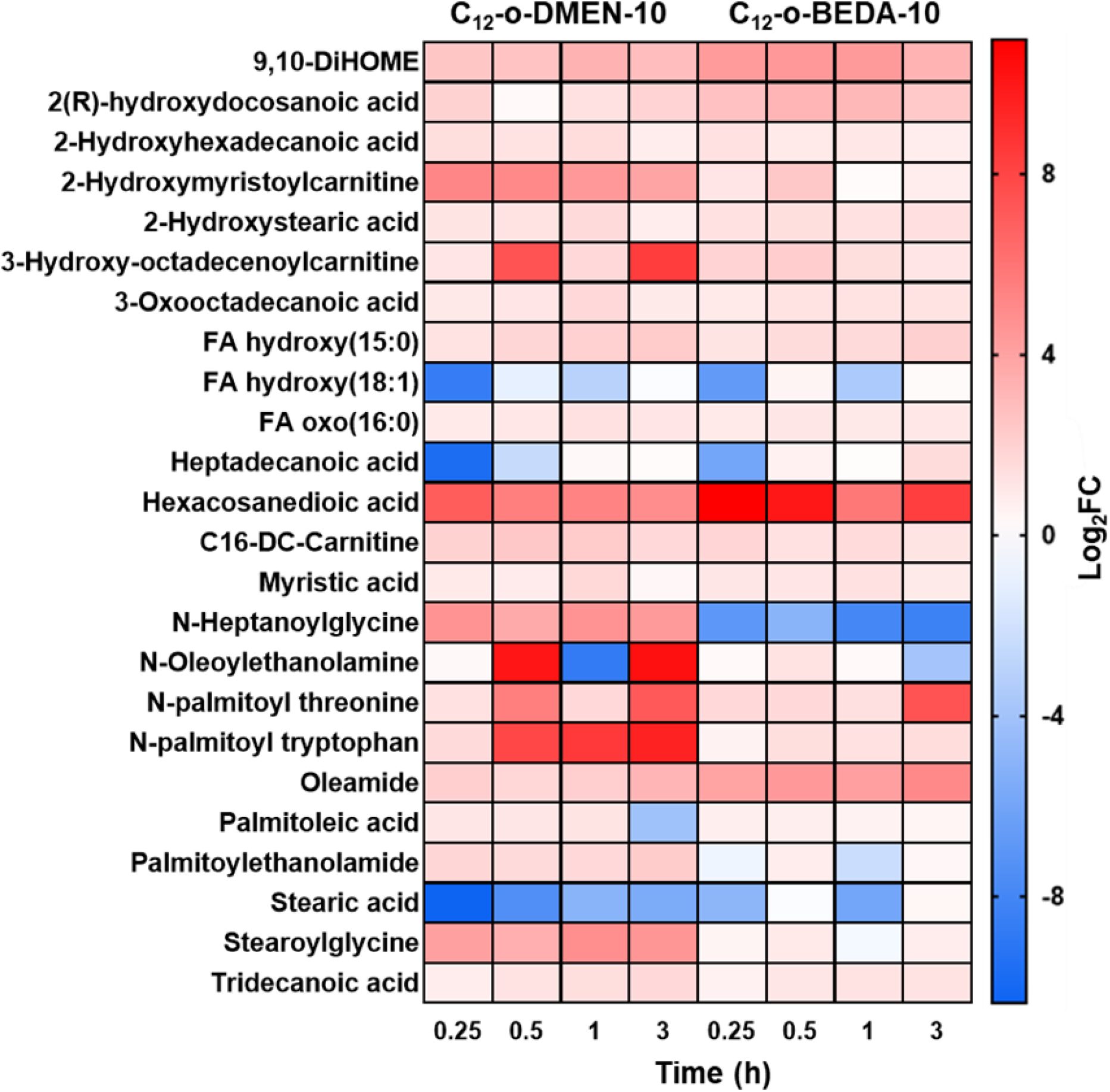
Heatmap representation of all significantly impacted metabolites (≥1.0-log_2_-FC; *p* < 0.05) involved in FA metabolism in MRSA ATCC 43300 after CLOs [C_12_-o-DMEN-10 and C_12_-o-BEDA-10] treatment at 0.25, 0.5, 1, and 3 h. Metabolites were putatively assigned based on accurate mass and predicted retention time (the position of the oxidation sites is proposed based on database entries, and were not experimentally confirmed

At 0.5 h, the perturbation of FA metabolism intensified, particularly under C_12_-o-DMEN-10 treatment. Notably, substantial elevations were observed in *N*-oleoylethanolamine, *N*-palmitoyl tryptophan, and hydroxy-octadecenoylcarnitine (≥ 7.0-log_2_FC, *p* < 0.05), indicative of enhanced FA conjugation and acyl-carnitine turnover [75]. In contrast, the corresponding increases induced by C_12_-o-BEDA-10 were markedly lower (≥ 1.0-log_2_FC, *p* < 0.05), suggesting a comparatively delayed and attenuated metabolic response. Both CLOs continued to promote elevated levels of hydroxylated FA, including hydroxyhexadecanoic acid and hydroxystearic acid (≥ 1.0-log_2_FC, *p* < 0.05), consistent with ongoing activation of lipid oxidation and cellular stress adaptation pathways [76] **(Fig. 4)**.

At 1 h, both CLOs induced persistent alterations in FA metabolism, with more pronounced effects observed under C_12_-o-DMEN-10. Notably, *N*-palmitoyl tryptophan, stearoylglycine, and *N*-heptanoylglycine remained significantly elevated (≥ 4.0-log_2_FC, *p* < 0.05), reflecting sustained activation of FA conjugation pathways. In contrast, these metabolites were suppressed or minimally altered by C_12_-o-BEDA-10. Both CLOs maintained increased levels of hydroxylated and oxidized intermediates such as oxooctadecanoic acid, hydroxyhexadecanoic acid, and FA oxo(16:0) (≥ 1.0-log_2_FC, *p* < 0.05), indicating ongoing FA-oxidation **(Fig. 4)**.

At 3 h, metabolic disruptions were sustained. Whereby the level of *N*-palmitoyl tryptophan, hydroxy-octadecenoylcarnitine, and *N*-palmitoyl threonine was significantly elevated (≥ 7.0-log_2_FC, *p* < 0.05), after C_12_-o-DMEN-10 treatment, indicating FA metabolic disruption. Oleamide and palmitoylethanolamide were also elevated (≥ 2.0-log_2_FC, *p* < 0.05). C_12_-o-BEDA-10 showed similar trends but less pronounced perturbations where the level of *N*-palmitoyl tryptophan, hydroxy-octadecenoylcarnitine, and *N*-palmitoyl threonine was elevated (≥ 1.0-log_2_FC, *p* < 0.05). Hydroxylated fatty acids, including hydroxydocosanoic acid and FA hydroxy(15:0), were modestly elevated (≥ 1.0-log_2_FC, p < 0.05) after both CLOs treatments. Meanwhile, stearic acid remained suppressed under C_12_-o-DMEN-10 but partially recovered under C_12_-o-BEDA-10, indicating distinct effects on saturated FA dynamics **(Fig. 4)**.

Taken together, both CLOs, induced time-dependent disruptions in FA metabolism, with C_12_-o-DMEN-10 consistently eliciting stronger and more sustained elevations in conjugated and hydroxylated fatty acid species compared to C_12_-o-BEDA-10.

## Conclusion

This study provides the first in-depth metabolomic investigation into the antimicrobial effects of two structurally distinct CLOs, C_12_-o-DMEN-10 and C_12_-o-BEDA-10, in *Staphylococcus aureus*, with a particular focus on glycerophospholipid and fatty acid metabolic pathways. Fluorescence-based membrane permeability assays (membrane disruption vs. growth inhibition using PI dye) had previously indicated a divergence in mechanisms of action between the two CLOs. However, the integration of untargeted metabolomics provided critical mechanistic clarity. Despite the minimal membrane permeability observed for C_12_-o-BEDA-10 by fluorescence-based assay, its metabolic signature closely mirrored that of C_12_-o-DMEN-10, marked by similar alterations in membrane lipid composition and fatty acid metabolism depicted by metabolomics. Both compounds induced time-dependent perturbations across multiple key membrane phospholipids, such as phosphatidylglycerol, the accumulation of lysophospholipids and phosphatidylethanolamine, and the remodelling of fatty acids. The elevated levels of hydroxylated and dicarboxylic fatty acids, as well as *N*-acyl amides and carnitine-linked intermediates, point to a broader stress adaptation response. These alterations collectively reflect a disruption in membrane-associated metabolic balance, resulting in compromised structural and functional membrane integrity. Importantly, both CLOs converged on similar metabolic outcomes, albeit with distinct temporal dynamics and magnitudes. These findings provide critical mechanistic insight into how CLOs exert their antibacterial activity and lay a strong foundation for further exploration of their potential as membrane-active therapeutic agents against drug-resistant Gram-positive pathogens. Also, the integration of metabolomics proved essential in revealing the underlying mechanisms of action, offering a comprehensive systems-level perspective that extends beyond the resolution of conventional qualitative assays. This study also highlights the value of metabolomic approaches in uncovering both direct and indirect modes of action, particularly when traditional methods such as PI fluorescence may not fully capture the extent or nature of bacterial response. Future work incorporating transcriptomic analyses will help clarify the regulatory drivers underpinning the metabolic shifts observed in this study. Furthermore, functional assays, such as genetic perturbation, enzyme activity tests, and metabolic rescue, will help distinguish primary CLO effects from secondary bacterial adaptations.

## Materials and Methods

### CLOs and antibiotic stock solution, media, and bacterial isolates

The CLOs [C_12_-o-DMEN-10 and C_12_-o-BEDA-10] were synthesised using Cu(0)-mediated reversible deactivation radical polymerization following the protocol established by Grace *et al*. [77]. The CLO stock solutions were formulated by initially dissolving the CLOs in DMSO, subsequently diluting with MilliQ water to achieve a 20% (v/v) DMSO concentration, and then vortexing until the solution was clear. DMSO was subjected to filtration using 0.22 µm sterile nylon filters before use. MRSA ATCC 43300, a genetically stable and globally recognised reference strain, was used to ensure reproducibility and minimise biological variability in the mechanistic assays performed in this study. While the use of a single strain enables experimental reproducibility and reduces biological variability in early-stage investigations, we recognise that strain-specific differences in resistance mechanisms may influence the generalizability of these findings. All susceptibility and time-kill experiments were conducted in cation-adjusted Mueller-Hinton broth (CAMHB; containing 20–25 mg/L Ca^2+^ and 10–12.5 mg/L Mg2+; BD, Sparks, MD, USA). Viable counting was conducted on cation-adjusted Mueller–Hinton agar (CAMHA; containing 25 mg/L Ca^2+^ and 12.5 mg/L Mg2+; BD, Sparks, MD, USA).

### Time Kill assay

A static concentration time-kill experiment of CLOs [C_12_-o-DMEN-10 and C_12_-o-BEDA-10] against MRSA ATCC 43300 was conducted (**Fig. S4**). Prior to the time-kill assay, MRSA ATCC 43300 was sub-cultured on a CAMHA plate and thereafter incubated at 37°C for approximately ∼18 to 24 hours. Three colonies were transferred from the CAMHA plate to inoculate 10 mL of sterile CAMHB in a 50 mL conical flask and incubated overnight in a shaking water bath at 37°C, 150 rpm for approximately 16 hours. The optical density (OD) of the bacterial suspension was quantified using a spectrophotometer, and the solution was suitably diluted to attain the desired initial inoculum of around ∼10^6^ CFU/mL. The inoculated flasks were treated with CLOs [C_12_-o-DMEN-10 and C_12_-o-BEDA-10] separately (in 20% DMSO) to attain concentrations of 128, 256, and 512 µg/mL for C_12_-o-DMEN-10 and 32, 64, 128, 256, and 512 µg/mL for C_12_-o-BEDA-10. The DMSO concentrations in the flaks were equal to or less than 0.125%. One culture flask served as a drug-free control. At 0, 0.5, 1, 3, 6, 24, and 48 hours, 1 mL samples were extracted from each flask, subjected to centrifugation, and rinsed twice with 0.9% normal saline. The samples were subsequently serially diluted in saline plates and inoculated onto CAMHA plates. Following a 24-hour incubation at 37°C, the colony-forming units (CFU) were enumerated, and the time-kill curves were plotted as log_10_ CFU/mL against time (hours).

### Metabolomics

#### Bacterial culture preparation

An untargeted metabolomics study was carried out to explore the mechanism(s) of action of CLOs [C_12_-o-DMEN-10 and C_12_-o-BEDA-10] separately against MRSA ATCC 43300 using a concentration of 256 µg/mL (i.e., 2-3× MIC). Samples were taken and analyzed at the 0.25-, 0.5-, 1-, and 3-h time points in four biological replicates. These early timepoints were chosen specifically to capture biochemical perturbations before extensive killing occurred. An overnight culture was prepared by inoculating a single colony into 100 mL CAMHB in 250 mL conical flasks (Pyrex) and incubating the suspension in a shaker at 37°C and 180 rpm for ∼16 h. After overnight incubation, log-phase cells were prepared in fresh MHB and then incubated for 2 h at 37°C at 180 rpm to the log phase with a starting bacterial inoculum of ∼10^8^ CFU/mL. Then, CLOs [C_12_-o-DMEN-10 and C_12_-o-BEDA-10] were separately added to obtain the desired concentration of 256 µg/mL (2-4× MIC), in parallel to a CLO-free control for each replicate. The flasks were then incubated at 37°C with shaking at 180 rpm. At each time point (0.25-, 0.5-, 1-, and 3-h), 15 mL samples were transferred to 50 mL Falcon tubes for quenching, and the OD reading at 600 nm (OD_600_) was then measured and normalized to the pre-treatment level of approximately ∼0.5 with fresh CAMHB. Samples were then centrifuged at 3,220 *× g* and 4°C for 10 min, and the supernatants were removed. The pellets were stored at −80°C until metabolite extraction. The experiment was performed in four biological replicates to reduce the bias from inherent random variation.

#### Metabolite extraction

The bacterial pellets were washed twice in 1 mL of 0.9% saline and then centrifuged at 3,220 *× g* and 4°C for 5 min to remove residual extracellular metabolites and medium components. The washed pellets were resuspended in a cold extraction solvent (chloroform-methanol-water at 1:3:1, v/v) containing 1 µM each of the internal standards 3-[(3-cholamidopropyl)-dimethylammonio]-1-propanesulfonate, *N*-cyclohexyl-3-aminopropanesulfonic acid, piperazine-N, N-bis (2-ethanesulfonic acid), and Tris. The samples were then frozen in liquid nitrogen, thawed on ice, and vortexed to release the intracellular metabolites (3x). Next, the samples were transferred to 1.5 mL Eppendorf tubes and centrifuged at 14,000 *× g* at 4°C for 10 min to remove any particulate matter. Finally, 200 µL of the supernatant was transferred into injection vials for liquid chromatography-mass spectrometry (LC-MS) analysis. An equal volume of each sample was combined and used as a QC sample. To ensure that the metabolite extraction specifically targeted live-stressed bacterial cells, we performed a series of optimization steps, including a high inoculum time-kill assay, as illustrated in **Fig 1**. By this method, we determined that there was at least 10^6.5^ (i.e., 3 million) CFU/mL present in the samples, which equates to >45 million viable CFU per harvested sample. Furthermore, it is crucial to highlight that the mechanism of action of the polymer, as previously tested and confirmed in this study, demonstrates a membrane-damaging effect. This characteristic supports the assertion that the percentage of dead but intact cells in the pelleted samples is expected to be low.

#### LC-MS analysis

Both hydrophilic interaction liquid chromatography (HILIC) and reversed-phase liquid chromatography (RPLC) coupled with high-resolution mass spectrometry were employed to ensure the detection of both hydrophilic and hydrophobic metabolites. Samples were analyzed on a Dionex U3000 high-performance liquid chromatography system in tandem with a Q-Exactive Orbitrap mass spectrometer (Thermo Fisher) in both positive and negative ion modes with a resolution of 35,000. The HILIC method was described previously in detail [78]. Briefly, samples maintained at 4°C were eluted through a ZIC-pHILIC column (5 µm, polymeric, 150 × 4.6 mm; SeQuant, Merck) by mobile phase A (20 mM ammonium carbonate) and mobile phase B (acetonitrile). The gradient started with 80% mobile phase B at a flow rate of 0.3 mL/min and was followed by a linear gradient to 50% mobile phase B over 15 min. The Ascentis Express C8 column (5 cm × 2.1 mm, 2.7 µm) (catalog no. 53,831-U; Sigma-Aldrich) was applied in the RPLC method. The samples were controlled at 4°C and eluted by mobile phase A (40% of isopropanol and 60% of Milli-Q water with 8 mM ammonium formate and 2 mM formic acid) and mobile phase B (98% of isopropanol and 2% of Milli-Q water with 8 mM ammonium formate and 2 mM formic acid). The linear gradient started from 100% mobile phase A to a final composition of 35% mobile phase A and 65% mobile phase B over 24 min at 0.2 mL/min. All samples were analyzed within a single LC-MS batch to avoid variations. The pooled quality control samples, internal standards, and total ion chromatograms were assessed to evaluate the chromatographic peaks, signal reproducibility, and stability of the analytes. To assist in the identification of metabolites, a mixture of ∼500 metabolite standards was analyzed within the same batch to confirm accurate mass and retention time.

### Data processing, bioinformatics, and statistical analyses

IDEOM was used to convert raw data obtained by LC-MS to perform semi-quantitative analysis of annotated metabolites [79]. ProteoWizard, a freely available software library for LC-MS data analysis, was first used to extract mzXML files from LC-MS raw data. These files were then processed using XCMS for peak picking [80]and MZmatch.R for peak alignment and filtering with minimum detectible intensity of 100,000 and RSD of 0.8. Using the same MZmatch.R tool, the missing peaks were also retrieved and annotated. Common sources of noise (contaminant signals, peak shoulders, and irreproducible peaks) were removed using the IDEOM workflow with default settings (version 23). The gain and loss of protons were corrected in positive and negative electrospray ionization mode and then a data-dependent mass recalibration (2 ppm) step for putative metabolites was performed. Metabolites confirmed with authentic standards were assigned with MSI level 1 identification. Putative metabolites (with MSI level 2 identification) were identified by comparing their accurate masses and retention times with the standards in the databases including KEGG (Kyoto Encyclopedia of Genes and Genomics), LipidMaps, MetaCyc, and HMDB. Peak height intensities were used for the quantification of the metabolites. Statistical analysis was performed using MetaboAnalyst 6.0 [81], a freely available online statistical tool. Briefly, putative metabolites were extracted from IDEOM and tabled per time point, then uploaded on MetaboAnalyst 6.0. Data were filtered using the interquartile range, normalized by the median, log_2_ transformed, and autoscaled. Fold change (FC) was calculated relative to the control from the corresponding time point. Univariate analysis was performed using two-sample t-tests (fold change threshold = 2; FDR adjusted *P*-value <0.05) to determine significantly perturbed metabolites for each time point. Multivariate analysis was performed and included the generation of heat maps and PCA plots. Finally, the KEGG IDs of metabolites were uploaded to KEGG Mapper [82], and pathways were constructed.

## Data availability statement

The data supporting this article have been included as part of the Supplementary Information. Further data will be made available on a reasonable request from reviewers.

## CRediT authorship contribution statement

**Muhammad Bilal Hassan Mahboob:** Conceptualization, Methodology, Software, Formal analysis, Investigation, writing – original draft, and editing.

**Jessica R. Tait:** Conceptualization, Methodology, Supervision, Validation, Writing – review & editing

**Dovile Anderson:** Resources, Data curation, Investigation, Software, Formal analysis

**Maytham Hussein:** Conceptualization, Methodology

**John F. Quinn:** Funding acquisition, Supervision, Validation, Writing – review & editing

**Tony Velkov:** Funding acquisition, Supervision, Validation

**Darren. J. Creek:** Resources, Supervision, Data curation, Validation, Writing – review & editing

**Michael R. Whittaker:** Conceptualization, Methodology, Funding acquisition, Supervision, Validation, Writing – review & editing

**Cornelia B. Landersdorfer:** Conceptualization, Methodology, Funding acquisition, Supervision, Validation, Writing – review & editing

## Declaration of competing interest

The authors declare that they have no known competing financial interests or personal relationships that could have appeared to influence the work reported in this paper.

## Acknowledgments

This work was supported by the Australian Research Council (DP200102829) and Monash Institute of Pharmacy and Pharmaceutical Sciences (MIPS). J. F. Q is grateful for an ARC Future Fellowship (FT170100144). We acknowledge the support of the Monash Proteomics and Metabolomics Platform.

## Notes

### Competing Interest Statement

The authors have declared no competing interest.

