## Supplemental material for "Bioactive Cationic Lipidated Oligomers (CLOs) as Antimicrobial Materials: Metabolomic Insights into MRSA Membrane Disruption"

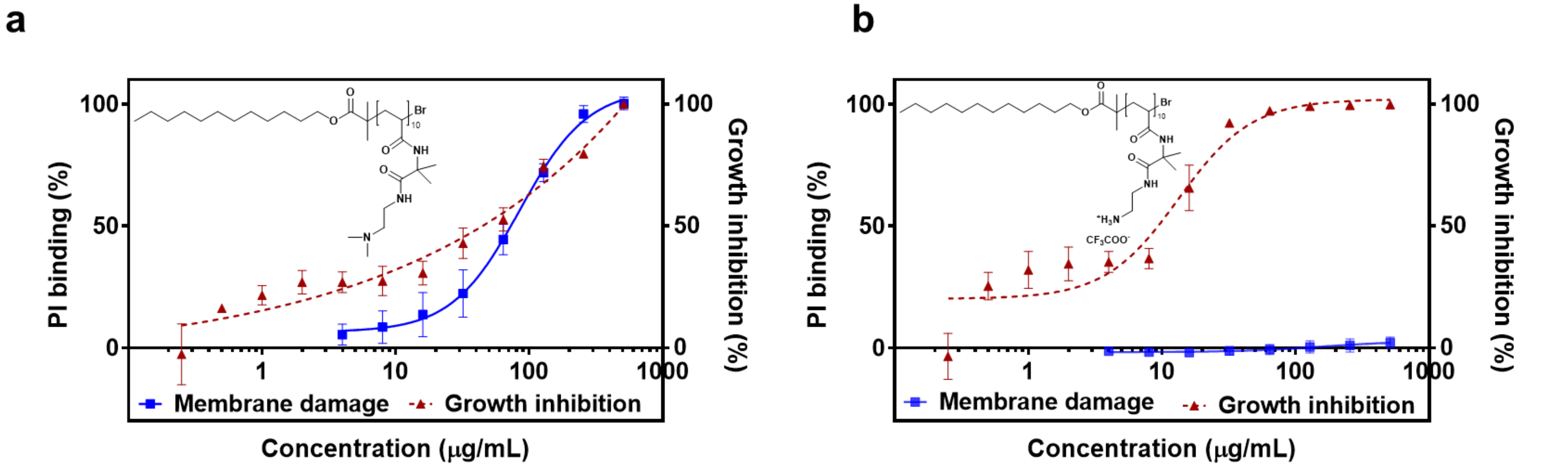


**Figure S1:** PI binding (%) and growth inhibition (%) assay of MRSA ATCC 43300 using **(a)** $C_{12}$-o-DMEN-10, **(b)** $C_{12}$-o-BEDA-10, over a range of oligomer concentrations. Data are presented as mean standard error of the mean (n = 6).

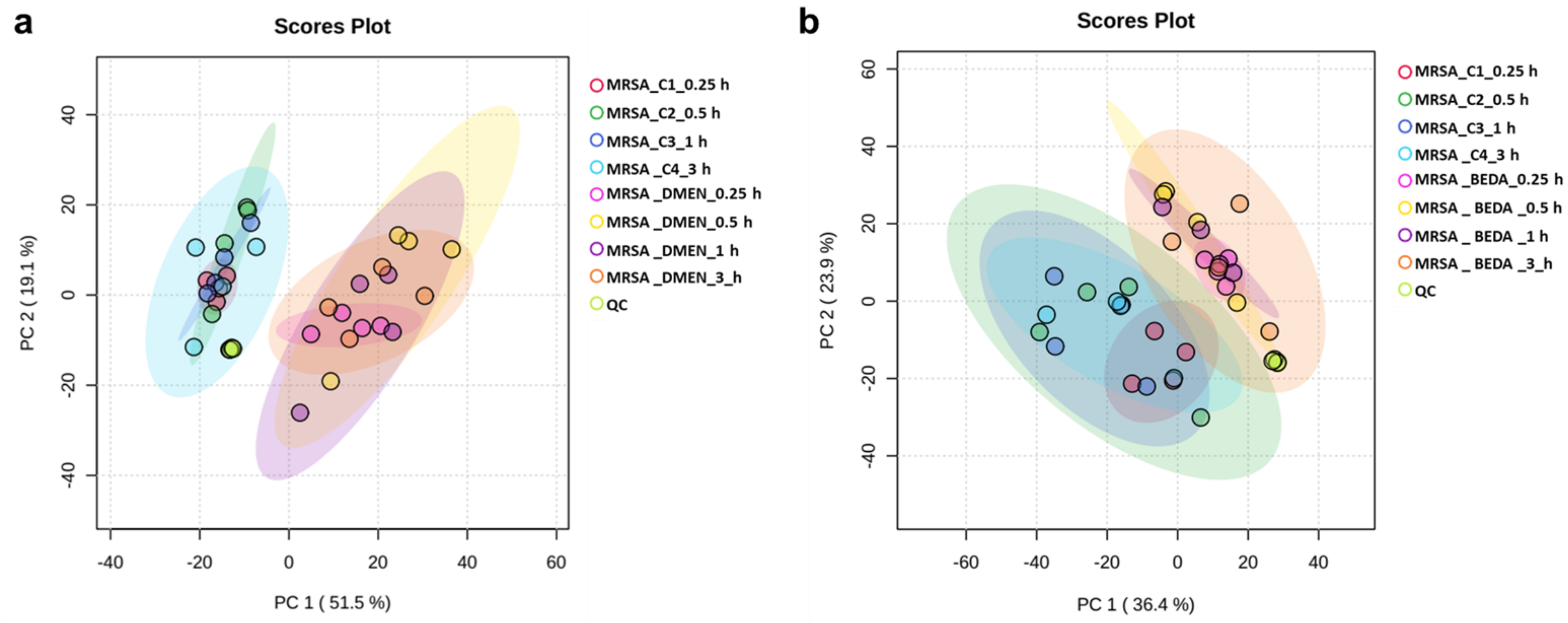


**Figure S2**. PCA plot of MRSA ATCC 43300 vs CLOs [$C_{12}$-o-DMEN-10 **(a)** and $C_{12}$-o-BEDA-10 **(b)**] at 0.25 h, 0.5 h, 1 h, and 3 h. DMEN= $C_{12}$-o-DMEN-10 and BEDA= $C_{12}$-o-BEDA-10.

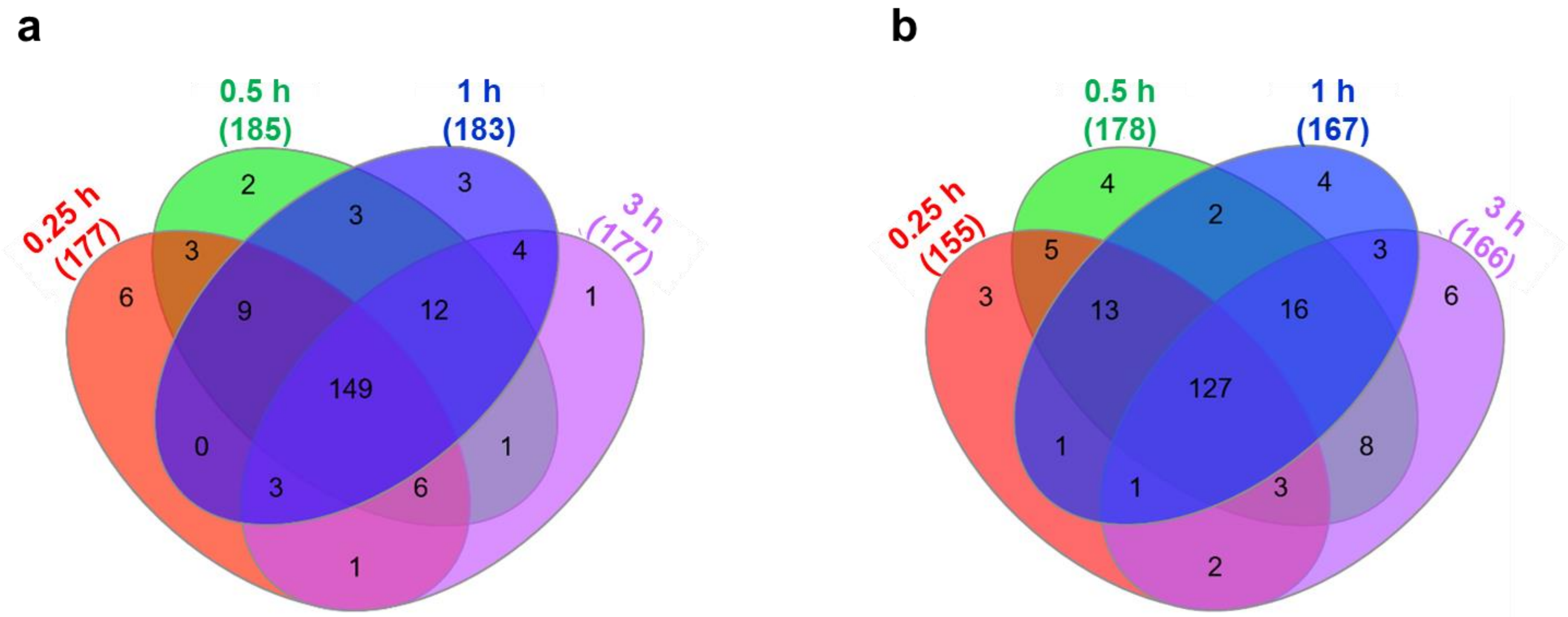


**Figure S3:** Venn diagram showing the number of metabolites of MRSA ATCC 43300, significantly affected by treatment with CLOs [$C_{12}$-o-DMEN-10 **(a)** and $C_{12}$-o-BEDA-10 **(b)**]. Significant metabolites were selected with ($\geq$1-$log_2$-FC; $p$<0.05).

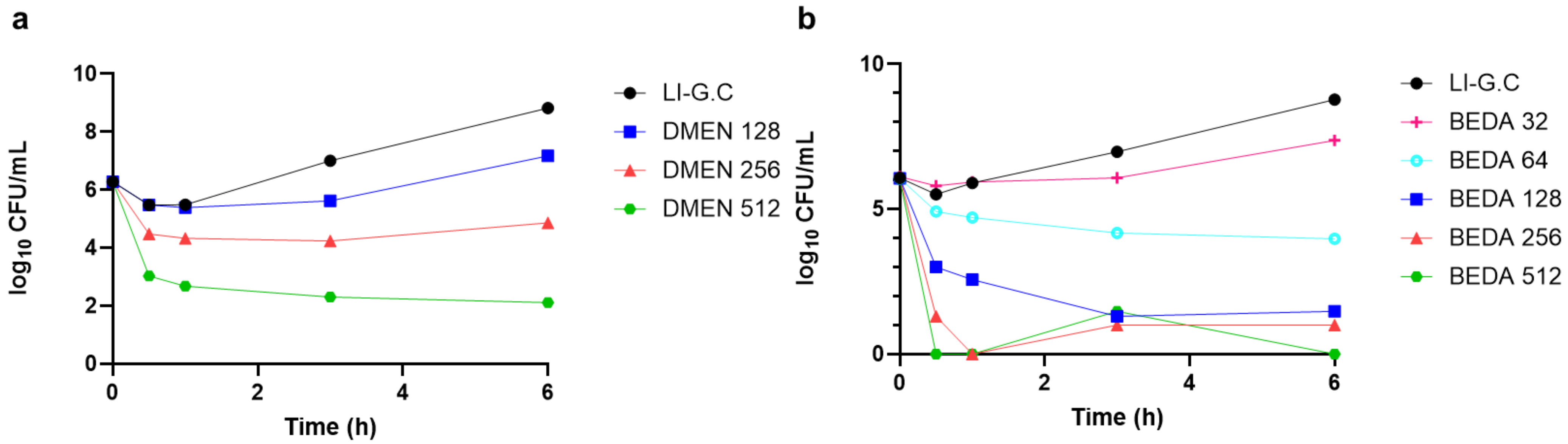


**Figure S4:** Killing kinetics of MRSA ATCC 43300 (initial inoculum ~$10^6$ $\log_{10}$ CFU/mL), after treatment with different concentrations of CLOs [**(a)** $C_{12}$-o-DMEN-10 (128, 256, 512 µg/mL) and **(b)** $C_{12}$-o-BEDA-10 (32, 64, 128, 256, 512 µg/mL)] (n=1).

**Table S1:** Significant metabolites putatively identified following exposure to CLOs [$C_{12}$-o-DMEN-10 and $C_{12}$-o-BEDA-10] in of methicillin-resistant *Staphylococcus aureus* (MRSA) ATCC 43300. Significant metabolites were determined using two-sample *t*-tests [($\log_2$fold-change [FC] $\geq$ 2, corresponding to a metabolite-level change of approximately 4-fold; false discovery rate [FDR] adjusted *p*-value $<$ 0.05)].

| Metabolomics changes ($Log_2FC$) | | | | | | | | | | |
|---|---|---|---|---|---|---|---|---|---|---|
| **Pathway** | **MAP** | **METABOLITES** | **$C_{12}$-o-DMEN-10** | | | | **$C_{12}$-o-BEDA-10** | | | |
| | | | **0.25 h** | **0.5 h** | **1 h** | **3 hr** | **0.25 h** | **0.5 h** | **1 h** | **3 hr** |
| **Lipids Metabolism** | **Lysophospholipids** | LysoPE(19:1) | 1.54 | 1.65 | 0.88 | 1.40 | 1.07 | 1.20 | 0.83 | 1.07 |
| | | LysoPE(14:0) | 0.83 | 1.35 | 0.75 | 0.05 | 0.47 | 0.62 | 0.99 | 0.33 |
| | | LysoPE(16:0) | -0.50 | -0.80 | -1.50 | -1.17 | 0.17 | 0.52 | 0.69 | 1.07 |
| | | LysoPE(16:1) | 0.71 | 0.61 | -0.31 | -0.29 | 1.94 | 1.60 | 1.23 | 1.50 |
| | | LysoPE(18:1) | 3.45 | 3.32 | 0.69 | 1.36 | 0.79 | 1.73 | 0.45 | 1.01 |
| | | LysoPG(16:0) | -0.36 | -0.01 | -0.18 | 0.14 | -0.86 | -1.27 | -0.30 | 0.17 |
| | | LysoPA(16:0) | -1.86 | -2.25 | -2.27 | -0.67 | -1.52 | -2.01 | -1.97 | -0.15 |
| | | LysoPG(17:0) | -1.34 | -1.36 | -1.52 | -1.15 | -1.51 | -1.77 | -1.05 | -1.01 |
| | **Glycerophospholipids** | PE(32:1) | 2.53 | 2.21 | 1.97 | 2.83 | 0.36 | -1.03 | 2.15 | 0.41 |
| | | PE(34:2) | -1.69 | -1.98 | 0.50 | -0.45 | -1.26 | -1.94 | -0.54 | -0.09 |
| | | PG (28:0) | -3.45 | -3.29 | -2.86 | -2.99 | -2.41 | -2.11 | -2.66 | -1.74 |
| | | PG(29:0) | -4.40 | -4.11 | -3.49 | -3.47 | -3.01 | -2.54 | -3.17 | -1.90 |
| | | PG(30:0) | -5.00 | -4.64 | -3.83 | -3.99 | -3.20 | -2.73 | -3.49 | -2.15 |
| | | PG(30:1) | -0.91 | -1.05 | -0.25 | -0.25 | -2.70 | -1.05 | -0.08 | 0.00 |
| | | PG(31:0) | -5.67 | -5.34 | -4.47 | -4.75 | -3.41 | -2.83 | -3.97 | -2.42 |
| | | PG(31:1) | -3.65 | -2.60 | -3.31 | -1.93 | -2.85 | -2.31 | -3.38 | -1.41 |
| | | PG(32:0) | -6.18 | -5.76 | -4.98 | -5.16 | -3.91 | -3.15 | -4.48 | -2.59 |
| | | PG(32:2) | -2.85 | 0.00 | 0.00 | 0.00 | -1.83 | 0.00 | 0.25 | 0.00 |

| | | | | | | | | | | |
|---|---|---|---|---|---|---|---|---|---|---|
| | | PG(33:0) | -6.52 | -5.84 | -5.07 | -3.14 | -4.06 | -3.12 | -4.79 | -0.81 |
| | | PG(33:1) | 0.25 | 0.25 | 1.15 | 0.25 | 0.90 | 0.77 | 0.70 | 0.25 |
| | | PG(34:0) | -7.12 | -6.24 | -5.53 | -5.30 | -4.57 | -3.60 | -5.29 | -2.78 |
| | | PG(34:1) | -4.41 | -3.67 | -2.93 | -2.28 | -4.32 | -2.46 | -4.06 | -2.77 |
| | | PG(34:2) | -1.21 | -0.25 | -0.25 | -1.40 | -1.21 | -0.25 | -0.13 | -1.10 |
| | | PG(35:0) | -7.04 | -6.44 | -5.45 | -5.53 | -5.00 | -3.68 | -5.76 | -3.81 |
| | | PG(35:1) | -5.86 | -4.10 | -4.79 | -1.77 | -4.26 | -2.88 | -4.88 | -2.58 |
| | | PG(35:2) | -2.02 | -0.76 | -1.66 | 1.76 | -4.75 | -1.29 | -2.81 | 0.00 |
| | | PG(36:0) | -5.14 | -3.21 | -3.24 | -4.85 | -5.47 | -3.21 | -5.57 | -3.22 |
| | | PI(22:0) | -1.31 | -1.05 | -0.95 | -2.04 | -0.33 | -0.72 | 0.23 | -0.50 |
| | | PS(32:0) | 0.74 | 0.25 | 0.62 | 0.80 | 2.07 | 2.60 | 2.05 | 1.39 |
| | | PS(34:0) | -1.35 | -0.94 | 0.06 | -0.12 | -1.36 | -0.42 | -0.80 | 0.06 |
| | **FA metabolism** | (9xi,10xi,12xi)-9,10-Dihydroxy-12-octadecenoic acid | 2.45 | 2.57 | 3.31 | 2.83 | 4.28 | 4.35 | 4.31 | 3.24 |
| | | 2(R)-hydroxydocosanoic acid | 1.90 | 0.25 | 1.26 | 1.88 | 2.66 | 3.16 | 3.03 | 2.29 |
| | | 2-Hydroxyhexadecanoic acid | 1.39 | 1.19 | 1.45 | 0.76 | 1.27 | 0.94 | 1.03 | 0.75 |
| | | 2-Hydroxymyristoylcarnitine | 5.20 | 5.05 | 4.37 | 3.90 | 1.11 | 2.35 | 0.15 | 0.78 |
| | | 2-Hydroxystearic acid | 1.13 | 1.21 | 1.57 | 0.78 | 1.26 | 1.39 | 1.26 | 1.36 |
| | | 3-Hydroxy-octadecenoylcarnitine | 1.10 | 7.39 | 1.63 | 8.35 | 1.86 | 2.13 | 1.41 | 1.11 |
| | | 3-Oxooctadecanoic acid | 0.97 | 1.12 | 1.62 | 0.87 | 0.91 | 1.24 | 1.20 | 1.17 |
| | | FA hydroxy(15:0) | 1.23 | 1.73 | 1.98 | 2.24 | 1.13 | 1.50 | 1.57 | 2.01 |
| | | FA hydroxy(18:1) | -1.13 | -0.14 | -0.39 | -0.03 | -0.88 | 0.47 | -0.46 | 0.17 |
| | | FA oxo(16:0) | 0.94 | 1.00 | 1.25 | 1.09 | 0.93 | 1.05 | 0.97 | 1.00 |
| | | Heptadecanoic acid | -1.27 | -0.32 | 0.30 | 0.14 | -0.79 | 0.56 | 0.09 | 1.49 |
| | | Hexacosanedioic acid | 7.01 | 5.51 | 5.34 | 4.88 | 10.98 | 10.03 | 5.81 | 8.26 |
| | | Hexadecanedioic acid mono-L-carnitine ester | 1.92 | 2.34 | 2.21 | 1.62 | 1.81 | 1.27 | 1.51 | 1.16 |
| | | Myristic acid | 0.93 | 0.83 | 1.66 | 0.32 | 1.06 | 1.08 | 1.29 | 0.90 |
| | | *N*-Heptanoylglycine | 4.61 | 3.62 | 4.65 | 4.33 | -0.91 | -0.66 | -1.04 | -1.09 |

| | | | | | | | | | | |
|---|---|---|---|---|---|---|---|---|---|---|
| | | *N*-Oleoylethanolamine | 0.28 | 10.07 | -1.15 | 10.27 | 0.25 | 1.20 | 0.25 | -0.51 |
| | | *N*-palmitoyl threonine | 1.29 | 5.54 | 1.63 | 7.13 | 1.62 | 1.62 | 1.31 | 7.37 |
| | | *N*-palmitoyl tryptophan | 1.57 | 7.89 | 8.52 | 9.51 | 0.55 | 1.38 | 1.26 | 1.45 |
| | | Oleamide | 2.06 | 1.71 | 2.07 | 3.18 | 3.96 | 4.38 | 4.09 | 5.07 |
| | | Palmitoleic acid | 1.03 | 1.06 | 1.14 | -0.53 | 0.71 | 0.69 | 0.52 | 0.42 |
| | | Palmitoylethanolamide | 1.75 | 1.55 | 1.61 | 2.16 | -0.09 | 0.81 | -0.29 | 0.30 |
| | | Stearic acid | -1.35 | -0.97 | -0.66 | -0.74 | -0.64 | -0.03 | -0.79 | 0.36 |
| | | Stearoylglycine | 4.05 | 3.44 | 4.82 | 4.48 | 0.44 | 0.94 | -0.06 | 0.77 |
| | | Tridecanoic acid | 0.77 | 1.23 | 1.40 | 1.60 | 0.58 | 1.06 | 1.20 | 1.16 |
| **Membrane Biosynthesis** | **Amino sugar and sugar nucleotides** | UDP-*N*-acetyl-D-galactosamine | -1.70 | -2.31 | -2.06 | -1.69 | -1.50 | -2.34 | -1.82 | -1.88 |
| | | *N*-Acetylneuraminate | -3.71 | -4.62 | -4.81 | -2.34 | -2.67 | -3.15 | -3.01 | -2.07 |
| | | D-Mannose 1-phosphate | -3.70 | -4.08 | -4.62 | -3.56 | -3.18 | -3.44 | -3.15 | -3.15 |
| | | UDP-glucose | -2.07 | -2.67 | -3.41 | -2.62 | -0.84 | -1.67 | -1.75 | -1.92 |
| | | 2-C-Methyl-D-erythritol 4-phosphate | -0.45 | -1.62 | -1.95 | -1.98 | -0.13 | -0.58 | -0.40 | -1.36 |
| | | alpha-D-Glucosamine 1-phosphate | -1.66 | -2.33 | -4.18 | -3.17 | -1.27 | -2.74 | -2.87 | -3.16 |
| | | *N*-Acetyl-D-glucosamine 6-phosphate | -1.01 | -2.67 | -1.12 | -1.49 | -0.50 | -2.02 | -1.25 | -1.80 |
| | **Peptidoglycan biosynthesis** | UDP-*N*-acetylmuramate | -1.60 | -2.24 | -2.73 | -1.64 | -0.07 | -0.75 | -1.44 | -0.76 |
| | | UDP-*N*-acetylmuramoyl-L-alanyl-D-glutamate | 2.91 | 0.13 | -3.00 | -3.38 | 3.16 | 0.01 | -1.26 | -1.50 |
| | | UDP-*N*-acetyl-2-amino-2-deoxy-D-glucuronate | -3.25 | -3.52 | -3.46 | -2.69 | -2.18 | -2.76 | -2.28 | -1.23 |
| | | D-Alanyl-D-alanine | -2.57 | -3.61 | -4.00 | -5.21 | -1.79 | -2.72 | -2.21 | -3.91 |
| | | UDPMurNAc(oyl-L-Ala-D-gamma-Glu-L-Lys-D-Ala-D-Ala) | -1.16 | -1.63 | -2.38 | -1.12 | 0.40 | -0.45 | -1.28 | -0.27 |
| | | L-Alanine | -1.67 | -2.78 | -2.57 | -2.90 | -1.11 | -2.48 | -1.15 | -2.66 |
| | | D-Glutamate | -2.71 | -3.71 | -3.96 | -2.99 | -1.90 | -2.92 | -2.55 | -2.73 |
| | | D-Aspartate | -3.06 | -3.88 | -4.26 | -3.29 | -2.26 | -3.29 | -2.88 | -3.44 |
| | **Teichoic Acid** | CDP-ribitol | -3.04 | -3.57 | -3.75 | -2.02 | -1.70 | -2.55 | -2.30 | -1.35 |
| | | CDP-glycerol | -2.85 | -2.84 | -4.10 | -2.61 | -1.58 | -2.08 | -2.39 | -1.83 |

| | | | | | | | | | | |
|---|---|---|---|---|---|---|---|---|---|---|
| | Lysine biosynthesis | D-Lysine | -1.89 | -3.17 | -3.66 | -1.24 | -1.33 | -2.69 | -2.50 | -1.09 |
| | | L-2-Aminoadipate | -2.31 | -2.37 | -2.45 | -3.55 | -2.00 | -2.03 | -2.12 | -3.52 |
| | | N6-Acetyl-L-lysine | -2.76 | -3.35 | -3.67 | -1.42 | -1.90 | -2.87 | -2.78 | -0.61 |
| | | Protein N6,N6-dimethyl-L-lysine | -3.20 | -2.87 | -3.06 | -3.07 | -1.87 | -2.59 | -2.66 | -3.20 |
| | | meso-2,6-Diaminoheptanedioate | 0.03 | -2.81 | -1.08 | -1.87 | -0.62 | -2.67 | -1.08 | -1.87 |
| | | (2R,4S)-2,4-Diaminopentanoate | -0.98 | -1.64 | -2.31 | -1.59 | -0.57 | -1.61 | -1.43 | -1.82 |
| | | 5-Acetamidopentanoate | -4.41 | -4.76 | -4.60 | -3.04 | -3.49 | -4.44 | -3.68 | -2.12 |
| | **Mevalonic acid Pathway** | farnesyl phosphate | 2.16 | 1.89 | 1.70 | 0.17 | 2.98 | 1.03 | 0.92 | 0.01 |
| | | Mevalonic acid-5P | -2.56 | -2.95 | -6.79 | -4.78 | -4.40 | -3.19 | -2.63 | -3.07 |
| **Amino acid metabolism** | | Glycine | -1.16 | -1.70 | -1.94 | -2.22 | -0.83 | -1.51 | -1.08 | -2.29 |
| | | 2-Aminoheptanoate | -3.74 | -4.81 | -5.30 | -6.82 | -2.99 | -4.41 | -5.06 | -5.24 |
| | | L-Aspartate 4-semialdehyde | -3.87 | -4.57 | -4.94 | -4.48 | -3.14 | -3.71 | -3.46 | -3.55 |
| | | L-Glutamine | -3.92 | -5.05 | -5.51 | -2.99 | -3.31 | -4.37 | -4.22 | -2.73 |
| | | L-Methionine | -1.56 | -2.24 | -1.94 | -1.35 | -1.01 | -1.71 | -0.86 | -1.29 |
| | | L-Valine | -2.50 | -3.20 | -3.57 | -0.18 | -1.68 | -2.93 | -2.33 | -0.15 |
| | | 4-Aminobutanoate | -2.00 | -2.47 | -1.92 | -3.84 | -2.10 | -3.14 | -2.64 | -3.44 |
| | | O-Phospho-L-homoserine | -1.16 | -1.79 | -2.04 | -3.25 | -0.52 | -1.04 | -1.06 | -2.61 |
| | | Iminoglycine | -1.92 | -1.85 | -3.44 | -2.91 | -1.77 | -2.65 | -1.66 | -2.37 |
| | | *N*-Formyl-L-methionine | -4.04 | -4.10 | -3.78 | -4.84 | -2.87 | -3.04 | -2.37 | -3.70 |
| | | L-Asparagine | -2.01 | -2.79 | -0.94 | -2.71 | -2.58 | -2.68 | -0.87 | -3.66 |
| | | *N*-Acetyl-L-aspartic acid | -3.86 | -5.73 | -5.60 | -8.50 | -2.81 | -4.11 | -3.85 | -6.65 |
| | | malyl alpha-D-glucosaminide | -2.80 | -3.88 | -4.47 | -3.93 | -1.63 | -2.65 | -2.91 | -2.36 |
| | | malyl N-acetyl-alpha-D-glucosaminide | -1.84 | -2.25 | -3.30 | -2.93 | -1.19 | -1.88 | -2.76 | -1.91 |
| | | Glutamate methylester | -4.11 | -4.86 | -5.38 | -5.33 | -3.28 | -3.90 | -3.68 | -3.71 |
| **DNA and RNA biosynthesis** | **Purines biosynthesis** | ATP | -2.84 | -4.52 | -4.95 | -3.14 | -2.63 | -3.86 | -3.85 | -2.82 |
| | | ADP | -4.01 | -4.82 | -4.48 | -4.08 | -3.07 | -3.54 | -3.19 | -3.62 |
| | | dADP | -3.08 | -4.42 | -2.70 | -3.85 | -2.43 | -3.30 | -3.11 | -4.86 |
| | | dAMP | -2.71 | -4.00 | -5.40 | -3.07 | -1.89 | -2.64 | -3.10 | -3.70 |
| | | Adenine | -2.07 | -2.08 | -2.62 | -1.78 | -1.06 | -1.39 | -1.08 | -1.06 |

| | | | | | | | | | | |
|---|---|---|---|---|---|---|---|---|---|---|
| | | Adenosine | -3.72 | -3.57 | -4.60 | -5.48 | -2.25 | -2.65 | -2.62 | -3.08 |
| | | Deoxyadenosine | -3.35 | -3.30 | -2.51 | -3.40 | -1.19 | -1.80 | -1.58 | -2.70 |
| | | GDP | -3.68 | -4.27 | -4.91 | -6.42 | -3.32 | -4.77 | -4.19 | -5.80 |
| | | GTP | -3.13 | -4.85 | -5.64 | -5.90 | -3.53 | -5.70 | -5.41 | -5.71 |
| | | GMP | -1.20 | -4.64 | -4.85 | -6.31 | -2.78 | -4.01 | -3.34 | -4.85 |
| | | dGDP | -2.71 | -4.24 | -4.69 | -2.86 | -2.61 | -3.66 | -3.46 | -2.45 |
| | | Guanine | 1.55 | 1.14 | -0.17 | -2.51 | 2.01 | 0.74 | 0.27 | -2.49 |
| | | 8-Hydroxyadenine | -4.16 | -3.55 | -1.44 | -1.92 | -2.01 | -3.62 | -3.12 | -4.05 |
| | | *N*-Formiminoglycine | -2.87 | -3.67 | -4.43 | -4.34 | -1.77 | -2.71 | -2.95 | -3.23 |
| | | 5'-Phosphoribosyl-N-formylglycinamide | -5.98 | -2.90 | -2.98 | -5.49 | -3.90 | -5.12 | -5.03 | -4.57 |
| | | 1-(5'-Phosphoribosyl)-5-amino-4-(N-succinocarboxamide)-imidazole | -3.27 | -5.61 | -3.48 | -3.35 | -3.96 | -5.81 | -5.12 | -4.51 |
| | **Pyrimidines biosynthesis** | CMP | -3.11 | -2.91 | -3.22 | -3.98 | -3.32 | -3.51 | -2.99 | -3.86 |
| | | CTP | -4.29 | -4.32 | -5.75 | -3.02 | -3.51 | -5.83 | -6.32 | 0.00 |
| | | CDP | -3.96 | -4.82 | -4.98 | -2.90 | -3.12 | -4.22 | -3.71 | -2.46 |
| | | dCMP | -2.63 | -2.69 | 0.00 | -0.25 | -2.78 | -2.84 | 0.00 | -0.25 |
| | | dCTP | -2.88 | -3.74 | -4.92 | -2.33 | -2.55 | -3.97 | -4.49 | -2.23 |
| | | UDP | -4.88 | -5.65 | -5.87 | -4.89 | -3.74 | -4.52 | -4.02 | -4.26 |
| | | UTP | -5.67 | -6.76 | -7.62 | -5.32 | -5.00 | -6.30 | -6.35 | -4.77 |
| | | UMP | -5.11 | -5.29 | -5.07 | -3.92 | -3.61 | -3.74 | -2.83 | -3.50 |
| | | Uracil | -1.63 | -1.38 | -2.03 | -1.33 | -0.88 | -1.13 | -1.07 | -1.35 |
| | | Thymine | -1.74 | -2.21 | -2.79 | -1.70 | -1.31 | -2.00 | -2.33 | -1.72 |
| | | dTDP | -3.76 | -4.68 | -5.30 | -4.64 | -3.13 | -3.13 | -2.65 | -3.30 |
| | | dTTP | -2.38 | -2.57 | -5.50 | -2.56 | -2.40 | -2.64 | -3.13 | -3.88 |
| | | Uridine | -0.90 | -1.38 | -1.45 | -1.39 | -1.19 | -1.59 | -1.08 | -1.34 |
| | | Orotate | -0.44 | -0.94 | -2.76 | -0.11 | 0.65 | -0.76 | -1.70 | 0.00 |
| | | (S)-Dihydroorotate | -2.77 | -4.19 | -5.57 | -2.74 | -1.76 | -4.14 | -4.78 | -1.75 |
| | | *N*-Carbamoyl-L-aspartate | -3.72 | -5.22 | -6.62 | -2.24 | -2.54 | -4.70 | -5.07 | -1.16 |
| | **Glycolysis** | Phosphoenolpyruvate | -2.81 | -4.37 | -4.62 | -4.67 | -2.75 | -3.57 | -4.03 | -3.29 |

| | | | | | | | | | | |
|---|---|---|---|---|---|---|---|---|---|---|
| | | D-Fructose 1,6-bisphosphate | -3.55 | -3.15 | -4.97 | -3.40 | -3.79 | -2.96 | -4.03 | -3.88 |
| | | D-Glyceraldehyde 3-phosphate | -3.81 | -3.47 | -2.54 | -3.86 | -3.29 | -4.35 | -4.72 | -4.46 |
| | | Glucose 6-phosphate | -2.21 | -2.61 | -3.29 | -2.65 | -1.41 | -1.81 | -1.95 | -2.29 |
| | | Mannitol | -3.29 | -3.57 | -3.20 | -3.72 | -2.55 | -2.46 | -2.62 | -4.59 |
| | | Serine | -0.65 | -1.73 | -2.49 | -0.33 | 0.01 | -1.24 | -0.86 | -0.12 |
| | | 3-Phospho-D-glycerate | -2.16 | -3.47 | -3.54 | -3.86 | -2.15 | -3.50 | -3.43 | -2.16 |
| | | D-Fructose | -0.99 | -2.07 | -2.69 | -2.06 | -0.33 | -1.53 | -1.26 | -1.87 |
| | | Hydroxypyruvate | -2.48 | -4.33 | -4.27 | -2.22 | -2.21 | -3.94 | -3.43 | -0.21 |
| | **Tricarboxylic Acid (TCA) cycle** | b-aminoisobutyric acid | -1.15 | -1.40 | -1.48 | -1.71 | -0.81 | -1.07 | -0.79 | -1.58 |
| | | Acetyl-CoA | -2.92 | -3.30 | -4.33 | -4.39 | -2.98 | -3.71 | -3.29 | -2.57 |
| | | Thiamin diphosphate | -3.05 | -3.64 | -3.52 | -2.41 | -2.36 | -2.86 | -2.55 | -1.67 |
| | | Propanoyl-CoA | 8.34 | 4.72 | 11.52 | 7.84 | 0.48 | -0.88 | 2.70 | -0.98 |
| | | 2-Oxoglutarate | -0.58 | -1.68 | -1.76 | -0.68 | -0.28 | -0.94 | -0.52 | -1.43 |
| | | CoA | -5.75 | -4.18 | -6.72 | -5.93 | -3.16 | -3.90 | -3.81 | -3.79 |
| | | (S)-Malate | -0.92 | -2.04 | -2.67 | -2.99 | -1.16 | -2.57 | -1.84 | -1.57 |
| | | (S)-2-Hydroxyglutarate | -2.87 | -3.34 | -3.22 | -4.47 | -2.12 | -2.53 | -2.05 | -4.31 |
| | | Citrate | -0.40 | -0.88 | -1.15 | -1.44 | -0.32 | -1.38 | -0.64 | -1.58 |
| | | 4-Oxoglutaramate | -1.51 | -1.88 | -2.47 | -1.95 | -0.86 | -1.57 | -1.51 | -1.80 |
| | **Pentose phosphate pathway (PPP)** | 6-Phospho-D-gluconate | -2.72 | -3.48 | -4.43 | -3.50 | -2.17 | -3.30 | -3.20 | -3.05 |
| | | Sedoheptulose 1,7-bisphosphate | -2.96 | -3.37 | -5.86 | -4.93 | -2.83 | -3.26 | -4.57 | -4.79 |
| | | D-Glucono-1,5-lactone 6-phosphate | -3.77 | -2.57 | -3.23 | -3.95 | -2.45 | -2.49 | -3.27 | -3.86 |
| | | 2-Deoxy-D-ribose 5-phosphate | -2.31 | -2.56 | -3.11 | -3.13 | -1.64 | -3.54 | -5.51 | -1.93 |
| | | Sedoheptulose | -3.45 | -3.34 | -4.91 | -4.19 | -2.17 | -2.61 | -4.10 | -4.62 |
| | | D-Glucono-1,5-lactone6-phosphate | -3.25 | -1.19 | -1.83 | -2.39 | -2.51 | -2.49 | -1.83 | -2.39 |
| | | D-Ribose | 0.60 | 0.56 | 0.11 | 0.25 | 1.02 | -0.01 | 0.67 | -0.03 |
| | | Deoxyribose | -3.79 | -3.51 | -2.59 | -4.29 | -3.14 | -3.28 | -1.63 | -4.61 |
| | | Sedoheptulose 7-phosphate | -1.78 | -3.07 | -2.94 | -4.74 | -2.16 | -2.79 | -2.75 | -3.26 |
| **Shikimic Acid Pathway Metabolism** | | (10aS)-10,10a-dihydrophenazine-1-carboxylate | 1.36 | 1.72 | 0.37 | 0.46 | 2.95 | 2.99 | 2.72 | 1.51 |

| | | | | | | | | | | |
|---|---|---|---|---|---|---|---|---|---|---|
| | | (1R*,3R*,3'S*)-1,2,3,4-Tetrahydro-1-(2-thio-3-pyrrolidinyl)-beta-carboline-3-carboxylic acid | -6.24 | -2.86 | -3.35 | -5.04 | -5.21 | -6.09 | -5.58 | -5.34 |
| | | 2-O-methyl cyclic 3-deoxy-D-arabino-heptulosonate 7-phosphate | -3.23 | -4.13 | -4.73 | -4.00 | -2.60 | -3.77 | -3.29 | -3.18 |
| | | alpha-(2,6-anhydro-3-deoxy-D-arabino-heptulopyranosid)onate 7-phosphate | -2.03 | -3.47 | -3.19 | -4.12 | -1.35 | -2.83 | -3.60 | -4.69 |
| | | 2-keto-4-hydroxy-5-phosphopentanoate | -3.38 | -4.93 | -5.24 | -6.62 | -1.95 | -2.58 | -2.72 | -4.57 |
| **Energy Metabolism** | **Arginine biosynthesis** | L-Ornithine | -3.57 | -5.58 | -5.54 | -3.95 | -2.37 | -4.17 | -3.14 | -1.97 |
| | | *N*-Acetylornithine | -3.52 | -5.23 | -5.22 | -4.07 | -2.42 | -3.61 | -3.17 | -1.67 |
| | | *N*-Acetyl-L-glutamate | -4.03 | -5.20 | -5.91 | -5.26 | -2.59 | -3.29 | -5.66 | -3.63 |
| | | 4-Guanidinobutanal | -0.97 | -0.16 | -0.10 | -1.29 | -0.26 | -1.19 | -0.90 | -1.52 |
| | | Arginine | -0.95 | -1.59 | -2.11 | -1.57 | -0.57 | -1.50 | -1.29 | -1.74 |
| | | N2-Succinyl-L-arginine | -2.36 | -3.08 | -3.44 | -1.62 | -1.23 | -2.84 | -2.28 | -1.29 |
| | | N2-Succinyl-L-ornithine | -2.16 | -3.05 | -0.25 | -2.45 | 0.00 | -2.03 | 0.00 | -2.81 |
| | | *N*-Succinyl-L-glutamate | -4.45 | -4.54 | -5.17 | -2.94 | -2.43 | -4.06 | -4.48 | -2.63 |
| | | L-Citrulline | -2.43 | -4.04 | -3.90 | -4.90 | -1.83 | -3.31 | -2.14 | -2.99 |
| | **Histidine biosynthesis** | *N*-Formimino-L-glutamate | -3.31 | -3.42 | -3.39 | -2.73 | -2.60 | -2.93 | -1.86 | -0.29 |
| | | *N*-Acetylhistidine | -2.95 | -3.53 | -3.27 | -2.18 | -1.70 | -2.17 | -1.67 | -0.62 |
| | | Histidine | -2.59 | -3.93 | -3.52 | -2.18 | -1.75 | -3.60 | -2.28 | -2.63 |
| | | Ergothioneine | -2.91 | -3.44 | -2.83 | -0.93 | -2.35 | -2.90 | -1.69 | -0.40 |
| | **Vitamins biosynthesis** | FAD | -0.93 | -1.17 | -1.28 | -1.57 | -0.73 | -1.07 | -0.76 | -1.97 |
| | | FMN | -0.86 | -2.80 | -2.07 | -4.06 | -0.86 | -1.61 | -1.23 | -3.57 |
| | | 5-Aminolevulinate | -1.89 | -2.35 | -2.52 | -5.04 | -1.39 | -1.65 | -1.94 | -4.65 |
| | | 2-(Hydroxymethyl)-3-(acetamidomethylene)succinate | -2.69 | -3.10 | -2.88 | -2.34 | -0.02 | -2.58 | -2.35 | -0.68 |
| | | Pyridoxine | -1.67 | -4.22 | -3.38 | -3.20 | -2.86 | -3.04 | -2.94 | -3.03 |
| | | Pantothenate | -1.10 | -1.78 | -1.91 | -2.35 | 0.09 | -1.31 | -1.11 | -1.40 |
| | | Pantetheine 4'-phosphate | -3.57 | -3.21 | -3.90 | -4.81 | -2.13 | -2.83 | -2.56 | -3.76 |

| | | | | | | | | | | |
|---|---|---|---|---|---|---|---|---|---|---|
| | | 4-Methyl-5-(2-phosphooxyethyl)thiazole | -2.32 | -3.16 | -3.53 | -2.60 | -1.55 | -2.48 | -2.45 | -3.74 |
| | | 6,7-Dimethyl-8-(D-ribityl)lumazine | -3.57 | -4.34 | -4.43 | -3.64 | -2.38 | -2.77 | -2.03 | -1.59 |
| **Stress and homeostasis Metabolism** | **Osmotic stress** | *N*-Acetylputrescine | -0.07 | -0.05 | -0.63 | 0.00 | -0.10 | -1.10 | -0.05 | 0.13 |
| | | Cyclic ADP-ribose | -2.30 | -3.19 | -1.68 | -3.15 | -1.81 | -2.68 | -2.35 | -1.98 |
| | | Carnitine | -3.64 | -4.63 | -5.30 | -6.92 | -3.06 | -4.34 | -4.97 | -5.24 |
| | | *N*-acetyl-L-glutaminyl-L-glutamine amide | -1.37 | -1.52 | -1.55 | -0.92 | -0.72 | -1.50 | -0.20 | -1.15 |
| | | lauryl sulfobetaine | 3.35 | 0.25 | 0.00 | 0.60 | 2.48 | 4.60 | 2.75 | 2.83 |
| | | 2-O-(beta-D-mannosyl)-bis(myo-inositol) 1,3'-phosphate | -2.40 | -2.15 | -2.20 | -2.38 | -1.76 | -1.93 | -1.03 | -1.59 |
| | | Feruloylputrescine | -1.64 | -1.24 | -1.47 | -0.98 | 0.00 | 0.00 | 0.46 | -0.54 |
| | | Choline sulfate | -2.37 | -2.58 | -2.48 | -1.72 | -1.39 | -2.28 | -1.44 | -0.59 |
| | | L-Proline | -0.37 | -1.00 | -0.67 | -1.98 | -0.34 | -0.77 | -0.10 | -1.18 |
| | | Trehalose 6-phosphate | -1.62 | -1.50 | -3.35 | -3.39 | -0.66 | -0.83 | -1.91 | -2.58 |
| | **Oxidative stress** | CoA-glutathione | -1.98 | -2.35 | -3.66 | -3.98 | -1.34 | -2.66 | -2.29 | -5.02 |
| | | *N*-hydroxy-N-succinylcadaverine | -1.35 | -1.15 | -1.83 | -1.38 | -0.12 | -0.88 | -0.46 | -1.14 |
| | | NADPH | -4.88 | -5.31 | -6.11 | -5.63 | -4.02 | -5.12 | -3.67 | -3.33 |
| | | NADP+ | -2.90 | -3.49 | -3.77 | -3.05 | -2.19 | -2.83 | -2.72 | -2.56 |
| | | Ophthalmic acid | -1.53 | -2.66 | -2.45 | -3.76 | -1.75 | -2.28 | -2.45 | -4.99 |
| | | N4-acetyl-N4-hydroxy-1-aminopropane | -2.34 | -3.54 | -3.44 | -4.28 | -1.68 | -3.17 | -1.51 | -2.55 |
| | | L-Cystathionine | -3.63 | -4.56 | -4.89 | -4.07 | -2.66 | -3.67 | -3.03 | -2.39 |
| | | O-Acetyl-L-serine | -2.95 | -3.26 | -3.34 | -2.12 | -2.18 | -2.52 | -2.01 | -0.71 |
| | **Redox stress** | Succinate semialdehyde | -3.63 | -3.91 | -5.08 | -1.26 | -2.90 | -4.08 | -4.23 | -5.01 |
| | | (R)-Lactate | -4.38 | -4.40 | -4.90 | -4.64 | -3.69 | -3.97 | -4.15 | -3.89 |
| | **Membrane lipid remodelling** | 2-Hydroxymyristoylcarnitine | 5.20 | 5.05 | 4.37 | 4.14 | 1.11 | 2.35 | 0.15 | 0.81 |
| | | HpOTrE | -0.29 | -0.58 | -1.40 | -2.65 | 0.14 | 0.47 | -0.04 | 0.15 |
| | | *N*-Gluconyl ethanolamine phosphate | 0.00 | 0.00 | -3.72 | -3.70 | 0.00 | 0.00 | -2.15 | -2.10 |

| | | | | | | | | | | |
|---|---|---|---|---|---|---|---|---|---|---|
| | **Quorum sensing** | *N*-(butyryl)-L-homoserine | -3.66 | -4.37 | -4.45 | -4.25 | -2.92 | -4.04 | -3.48 | -3.05 |
| | | P-DPD | -3.54 | -4.41 | -5.24 | -6.62 | -1.95 | -2.58 | -2.72 | -4.57 |
| | **Misc.** | dTDP-D-desosamine | -3.09 | -3.88 | -5.18 | -1.59 | -2.66 | -3.89 | -4.06 | -5.64 |
| | | mycaminose | -2.72 | -3.32 | -5.02 | -7.03 | -2.31 | -2.72 | -4.33 | -6.42 |
